# Photoperiodic Flowering Response of Essential Oil, Grain, and Fiber Hemp (*Cannabis sativa* L.) Cultivars

**DOI:** 10.1101/2021.05.13.444025

**Authors:** Mengzi Zhang, Steven L. Anderson, Zachary Brym, Brian J. Pearson

## Abstract

Cultivation of hemp (*Cannabis sativa* L.) in tropical and subtropical regions can be challenging if flowering behavior of a given cultivar is unknown, poorly understood, or not accurately selected for the photoperiod. Identifying cultivars adapted to local environmental conditions is key to optimizing hemp vegetative and flowering performance. We investigated the effects of varying light cycles in regulating extension growth and flowering response of 15 essential oil and 12 fiber/grain hemp cultivars both in indoors and outdoors. Plants were subjected to eleven photoperiods in the controlled rooms ranging from 12 h to 18 h, and natural day length in the field. The critical photoperiod threshold was identified for seven essential oil cultivars and two fiber/grain cultivars. ‘Cherry Wine-CC’, ‘PUMA-3’, and ‘PUMA-4’ had the shortest critical day length between 13 h 45 min and 14 h. The flowering of essential oil cultivars was generally delayed by 1 to 2 d when photoperiod exceeded 13 h compared to 12 h, and flowering was further delayed by 7 to 8 d when photoperiod exceed 14 h. In fiber/grain cultivars, flowering was generally delayed by 1 to 3 d when day length exceeded 14 h. Flowering for most essential oil cultivars was delayed by 5 to 13 d under 14 h photoperiod compared to 13 h 45 min, suggesting a photoperiod difference as little as 15 min can significantly influence the floral initiation of some essential oil cultivars. Cultivars represented by the same name but acquired from different sources can perform differently under the same environmental conditions, suggesting genetic variation among cultivars with the same name. Average days to flower of fiber/grain cultivars was correlated with reported cultivar origin with faster flowering occurring among northern cultivars when compared to southern cultivars. Plant height generally increased as the day length increased in essential oil cultivars but was not affected in fiber/grain cultivars. In addition, civil twilight of approximately 2 µmol·m^-2^·s^-1^ was discovered to be biologically effective in regulating hemp flowering. Collectively, we conclude that most of the essential oil cultivars and some southern fiber/grain cultivars tested express suitable photoperiods for tropical and sub-tropical region cultivation.

## INTRODUCTION

Cultivation of hemp (*Cannabis sativa* L.) within the U.S. was restricted in 1937 following passage of the Marihuana Tax Act. Similarly, hemp cultivation was prohibited throughout the western world during most of the 20th century (Cherney and Small, 2016; Congressional Research Service, 2019). With the legal status of *Cannabis* production shifting in the U.S. following passage of the 2014 and 2018 farm bills (Agricultural Act of 2014, P.L. 113-79; Agriculture Improvement Act of 2018, P.L. 115-334), restrictions on hemp production were relaxed and interest in hemp cultivation thereafter rapidly increased. Within the U.S., classification of *Cannabis* is based upon the concentration of Δ9-tetrahydrocannabinol (THC) present in plant tissue. Plants with a concentration of ≤ 0.3% THC on a dry weight basis are legally recognized as industrial hemp whereas plants containing > 0.3% are recognized as marijuana, a Schedule I drug as defined by the Controlled Substances Act of 1967 (Congressional Research Service, 2019). Industrial hemp is commercially cultivated for its fiber, seed, and secondary metabolites. Hemp is used to produce a wide variety of industrial and consumer products to include food and beverages, personal care products, nutritional supplements, therapeutic products, fabrics and paper, and construction materials (Congressional Research Service, 2019). In 2016, the global fiber hemp market was valued at nearly $700 million with an expected growth rate of 10-20%, whereas the hemp-derived cannabidiol (CBD) market in 2022 is expected to be more than two-fold greater than it was in 2018 to become a $1.3 billion-dollar market (Anderson et al., 2019; Hemp Business Journal, 2018).

Hemp can be challenging to cultivate in tropical and subtropical regions given high temperatures, high humidity, and ample presence of disease and pests. Relatively short daylengths experienced in tropical and subtropical environments, however, arguably present the greatest challenge to the successful cultivation of hemp at lower latitudes. Hemp is considered an annual, dioecious, short-day plant (SDP) originating from temperate regions of Central Asia. Most hemp varieties are photoperiodic and thus flowering of hemp is dependent upon day length or photoperiod. *Cannabis* has adapted to a wide range of climates and latitudes (23 to 52°N) and thus can possess large variability in its sensitivity to day length (Zhang et al., 2018). Timing of transition from vegetative growth to flowering is key for high yield and acceptable fiber quality of hemp (Amaducci et al., 2012). Earlier seasonal planting under critical daylength can extend the vegetative growth period prior to late-summer flowering, which is expected to occur generally 4 to 5 weeks after summer solstice in the northern hemisphere, dependent upon hemp variety and latitude (Anderson et al., 2019; Cherney and Small, 2016). Relatively short day length experienced in tropical and subtropical regions result in reduced vegetative growth and early seasonal transition to flowering that ultimately limits stem elongation and fiber biomass yield, key factors for successful commercial cultivation of industrial hemp (Cosentino et al., 2012; Hall et al., 2014). Thus, genotypes of hemp adapted to higher latitudes would be expected to perform poorly when cultivated in tropical and subtropical environments due to premature flowering and the negative influence it has on plant growth and yield (Amaducci et al., 2008; Cosentino et al., 2012; Hall et al., 2012).

Hemp expresses broad genetic diversity in hemp photoperiod requirements for vegetative-to-reproductive transition requirements, similar to that seen in other major crops (e.g. maize; Navarro, et al., 2017). Identifying plant genotypes adapted to a region’s light conditions is key to the successful cultivation of photoperiod crops, such as hemp (Cho et al., 2017; Jung and Müller, 2009). Zhang et al. (2018) discovered *Cannabis* can be generally categorized in three northern hemispheric haplogroups distinguished by geographical location (north of 40°N, 30 to 40°N, and south of 30°N); however, a myriad of photoperiod responses can be observed when breeding among haplogroups. Hemp selected for fiber production is generally believed to be a quantitative SDP with a relatively long photoperiod, around 14 h (dependent upon origin of the plant material). Excluding European varieties, the photoperiod response of most industrial hemp is poorly documented. ‘Kompolti’ (Hungarian variety) and ‘Futura 77’ (French variety) have an estimated maximal optimum photoperiod of 13.8 and 14 h, respectively (Heslop-Harrison and Heslop-Harrison, 1969). An estimated photoperiod of roughly 14 h was identified by Amaducci et al. (2008) for five European hemp cultivars as the most important single factor controlling flowering date. Flowering was increasingly delayed at longer photoperiods, but a 24 h photoperiod did not prevent ‘Fedrina 74’ (French variety) and ‘Kompolti Hybrid TC’ (Hungarian variety) from flowering (Lisson et al., 2000; Van der Werf et al., 1994). A Portuguese fiber variety was reported to have a maximum optimal photoperiod of 9 h, although the critical photoperiod is somewhere between 20 and 24 h (Heslop-Harrison and Heslop-Harrison, 1972; Lisson et al., 2000). In contrast to European hemp, flowering of Chilean and Kentucky hemp varieties occurred promptly under a photoperiod of 14 h or fewer but was considerably delayed or failed to flower when photoperiod exceeded 16 hours (Borthwick and Scully, 1954). A subtropical Australian variety, ‘BundyGem’, had a critical photoperiod between 13 h 40 min and 14 h 40 min, and plant maturity was significantly delayed when day length exceeded 14 h 40 min (Hall et al., 2014). An 11 to 12 h photoperiod has been reported to induce flowering of Thai hemp (Sengloung et al., 2009). While most of the studies on hemp was conducted in the field or greenhouses, plant responses to environment factors in a more strictly controlled environment, such as growth chambers, are very limited.

Until recently, *Cannabis* plants grown for recreational use have largely been cultivated indoors using artificial lighting. Given most of these cultivation operations were conducted prior to legalization of marijuana and were thus illegal, critical photoperiod of these types of *Cannabis* plants are not documented in the literature and information is limited. A day length of 12 h and 18 h are common practices to induce flowering or keep plants vegetative, respectively (Potter, 2014). Moher et al. (2020) indicated that *C. sativa* ‘802’, which although is not categorized as hemp given its 15 to 20% THC content, had a critical photoperiod between 15 to 16 hours. Growth chamber environments are ideal for investigating photoperiodism of hemp. With artificial lighting (typically from light-emitting diodes) being the only radiation source indoors, photoperiod is strictly controlled by the hours of light operation. In tropical and subtropical regions, vegetation of hemp under long days can be achieved in protected environments, such as greenhouses, by manipulating photoperiod utilizing end-of-day extension lighting and night interruption techniques that have been utilized in production of other common SDPs (Lane et al., 1963; Vince-Prue and Canham, 1983; Runkle et al., 1998; Zhang and Runkle, 2019). However, since hemp is often cultivated outdoors to reduce production costs, it is imperative that it is germinated or transplanted at timing with respect to natural photoperiod. Prediction of flowering time in response to a specific, known photoperiod is thus critical to support successful production both outdoors and indoors and optimization of select hemp varieties for a diverse range of growing regions. To directly address these needs, we utilized 7 growth chambers and investigated 15 cultivars of essential oil hemp and 12 cultivars of fiber/grain hemp to (i) empirically define critical photoperiod thresholds to induce vegetative to floral transition in diverse hemp cultivars, (ii) compare critical photoperiod thresholds to flowering dates within a subtropical field environment, and (iii) quantify physiological response of hemp cultivars under different photoperiod treatments.

## MATERIALS AND METHODS

### Expt.1: Photoperiod trial for selected essential oil cultivars

#### Seedling preparation and vegetative stage

Seeds, cuttings, or plants of all essential oil cultivars were obtained from three different sources (Supplementary table 1). Cultivars were selected based upon commercial interest and availability. Seeds of five essential oil cultivars, ‘Cherry Wine-BS’, ‘Cherry Blossom-BS’, ‘Cherry*T1-BS’, ‘Berry Blossom-BS’, and ‘Cherry Blossom-Tuan-BS’, were sown in 72 round cell propagation sheets (DPS72, The HC Companies, Twinsburg, OH) within Pro-Mix soilless substrate (HP Mycorrhizae Pro-Mix; Premier Tech Horticulture Ltd., Quakertown, PA, United States) containing 65-75% peat, 8-35% perlite, dolomite limestone, and mycorrhizae on November 19, 2019. Cuttings of the other 10 essential oil cultivars, ‘ACDC-AC’, ‘Super CBD-AC’, ‘Cherry-AC’, ‘Wife-AC’, ‘Cherry Blossom-BC’, ‘JL Baux-CC’, ‘ACDC-CC’, ‘Cherry Wine-CC’, ‘Cherry-CC’, and ‘Wife-CC’, were propagated on November 25, 2019. Each cultivar was propagated from identical mother stock plants to reduce potential genetic diversity among replicates. Stems of plant propagules were dipped into rooting hormone (Dip’N Grow; Dip’N Grow Inc., Clackamas, OR, United States) containing 1000 ppm indole-3-butyric acid (IBA) / 500 ppm napthaleneacetic acid (NAA) and then inserted into 3.8 cm rockwool cubes (Grodan; ROXUL Inc., Milton, Canada) that were pre-soaked with water containing a pH of 5.8 as per manufacturer recommendations. Both seeded trays and cuttings were grown at 25 □ under 24 h photoperiod and were hand irrigated daily as needed.

After roots were well established (∼21 d), the most uniform rooted propagules of each cultivar were selected and transplanted into 1.1 L containers (SVD-450, T.O. Plastics, Clearwater, MN, U.S.) filled with Pro-Mix soilless substrate and top-dressed with 5 g of Osmocote Plus 15-9-12 5-6 month slow-release fertilizer (Everris NA, Inc.; Dublin, OH, United States) containing 7% ammoniacal and 8% nitrate nitrogen, 9% phosphate and 12% soluble potash on December 17, 2019. Plants were randomly assigned to seven identical controlled rooms with 10 replicates per cultivar in each room and were cultivated at 25 □ under a photoperiod of 18 hours (0600 HR to 2400 HR) for vegetative growth. Plants were irrigated for 4 min every 5 days for the first two weeks and 4 min every 3 days thereafter as controlled by an automatic irrigation system.

#### Lighting treatments during flowering stage

After three weeks of vegetative growth following transplant, seven lighting treatments were randomly assigned to each controlled room on January 7, 2020. Ten plants of each hemp cultivar were grown at 25 □ under the photoperiod of 12 h (0600 HR to 1800 HR), 12 h 30 min (0600 HR to 1830 HR), 13 h (0600 HR to 1900 HR), 13 h 30 min (0600 HR to 1930 HR), 13 h 45 min (0600 HR to 1945 HR), 14 h (0600 HR to 2000 HR), and 18 h (0600 HR to 2400 HR) provided by light-emitting diodes (LEDs) (VYPR 2p; Fluence Bioengineering, Inc., Austin, TX, United States). Lighting treatments were maintained until termination of the experiment five weeks following the vegetative growth period.

#### Environmental conditions during seedling, vegetative, and flowering stage

Propagation of cuttings and germination of seeds was conducted indoors in an environmentally controlled propagation room at the Mid-Florida Research and Education Center (Apopka, FL, United States). Air temperatures were maintained in all indoor grow rooms utilizing air conditioners set to 25 □. Air temperature and relative humidity data was collected every 10 minutes by thermocouples installed at plant canopy height and data was recorded utilizing a wireless data logging station (HOBO RX3000; Onset Computer Corporation, Bourne, MA, United States) every 10 minutes. The average temperature in the propagation room was 24.9 ± 0.04 □. Within the propagation room, 24 h photoperiod was provided by fluorescent lamps (E-conolight; Sturtevant, WI, United States) as sole-source lighting. The photosynthetic photon flux density (PPFD) on the propagation bench was measured by a quantum sensor (MQ-500; Apogee Instruments Inc., Logan, UT, United States) at 10 representative positions at the seedling canopy level. Average PPFD that cuttings and seedlings received was 53.9 ± 3.02 and 73.5 ± 3.61 µmol·m^-2^·s^-1^, respectively, with a daily light integral of approximately 4.7 and 6.4 mol·m^-2^·d^-1^, respectively.

Following transplant, all plants were cultivated in seven identical environmentally controlled rooms. Each room was equipped with two sole-source LEDs (VYPR 2p; Fluence Bioengineering, Inc., Austin, TX, United States) regulated by a timer (Titan Controls Apollo 8; Hawthorne Gardening Company, Vancouver, WA, United States) to provide varying controlled photoperiod treatments. A PPFD of approximately 300 and 330 µmol·m^-2^·s^-1^ was maintained at plant canopy height at the onset of the vegetative stage and flowering stage, respectively. Average temperature, relative humidity, and light intensity for vegetative and flowering stages for each lighting treatment are reported in Supplementary table 2.

#### Plant measurements and data collection

Flowering of female hemp plants is defined as the appearance of dual, fork-shaped stigmas protruding from tubular bracts (Hall et al., 2012) being visible at the apical meristem or decimal code of 2201 defined by Mediavilla et al., (1998). Flowering of male hemp plants is defined when five radial segments of the first pointed male bud open and start to release pollen (Hall et al., 2012) or decimal code of 2101 (Mediavilla et al., 1998). Plant height (from the substrate surface to the tallest meristem) was measured at the initiation of lighting treatments and at flowering. Extension growth was calculated by subtracting initial plant height from height at flowering. Days to flower and plant sex were recorded when plants initiated flowering. Boolean evaluation of plant flowering status was conducted at the end of week 5 following initiation of lighting treatments.

#### Experimental design and data analysis

The experiment was conducted using a complete randomized design with seven lighting treatments and multiple replicates. Each plant was considered an experimental unit. Data were pooled from multiple replicates and were analyzed with a restricted maximum likelihood mixed model analysis in JMP^®^ Pro 15 (SAS Institute, Inc., Cary, NC, United States) with post-hoc mean separation tests performed using Tukey’s honest significant difference test at *P* ≤ 0.05.

### Expt.2: Photoperiod trial for fiber and grain cultivars

#### Seedling preparation and vegetative stage

Twelve fiber/grain hemp cultivars from six different source origins were purchased, including Canadian cultivars - ‘CFX-1’ and ‘Joey’; Polish cultivar - ‘Tygra’; Southern European cultivars - ‘Carmagnola Selezionata’, ‘Helena’, ‘Fibranova’, and ‘Eletta Campana’; Northern Chinese cultivar - ‘HAN-FN-H’; Central Chinese cultivars - ‘HAN-NE’ and ‘HAN-NW’; and Southern Chinese cultivars - ‘PUMA-3’ and ‘PUMA-4’. Seeds were sown in a 72 round cell propagation sheets (DPS72, The HC Companies, Twinsburg, OH) filled with Pro-Mix HP soilless substrate on February 18, 2020. They were placed under a mist bench in a greenhouse and grown at 25 □ under natural daylight supplemented with 1000 W metal halide lighting to maintain an 18 h photoperiod. Seedlings were misted for 1 min at 8 am, 12 pm, and 5 pm each day.

Seedlings possessing the most uniform height were selected three weeks after germination when roots were well established and transplanted into 1.1 L containers, as described above, with Fafard 4P potting media (Sun Gro Horticulture Canada Ltd., Agawam, MA, United States) containing 48% peat, 30% pine bark, 10% perlite, and 12% vermiculite, top-dressed with 5 g of Osmocote Plus slow-release fertilizer as described above. Plants were randomly assigned to seven identical environmentally controlled rooms with 10 replicates for ‘CFX-1’, ‘Tygra’, ‘Helena’, ‘Eletta Campana’, ‘HAN-FN-H’, and ‘HAN-NE’; 9 replicates for ‘PUMA-3’; 7 replicates for ‘Joey’ and ‘Fibranova’; replicates for ‘PUMA-4’; 4 replicates for ‘Carmagnola Selezionata’ and ‘HAN-NW’ due to poor germination rates. Plants were grown at 25 □ under 18 h photoperiod (0600 HR to 2400 HR) for vegetative growth until initiation of photoperiod treatments.

#### Lighting treatments during flowering stage

Seven lighting treatments were randomly assigned to each controlled room after two weeks of plant vegetative growth on March 23, 2020. Twelve fiber/grain hemp cultivars were subjected to seven photoperiod treatments: 12 h (0600 HR to 1800 HR), 13 h 30 min (0600 HR to 1930 HR), 13 h 45 min (0600 HR to 1945 HR), 14 h (0600 HR to 2000 HR), 14 h 30 min (0600 HR to 2030 HR), 14 h 45 min (0600 HR to 2045 HR), and 18 h (0600 HR to 2400 HR) provided by LEDs. Treatments were selected based on the common photoperiod range of fiber/grain cultivars documented in literature and the expected photoperiod of tropical and subtropical regions. Lighting treatments were maintained for five weeks before the termination of the experiment.

#### Environmental conditions during seedling, vegetative, and flowering stage

Germination of seedlings was conducted in a research greenhouse under a mist bench. Greenhouse heaters and fans were controlled by an environmental control system (Wadsworth Control System, Arvada, CO, United States) and set to operate when greenhouse temperature was ≤ 16 □ or ≥ 24 □, respectively. Seedlings were misted for a duration of 1 min three times per day utilizing a programmable irrigation controller (Sterling 12; Superior Controls Co., Inc., Valencia, CA, United States) and subjected to an 18 h photoperiod (from 0700 HR to 0100 HR) with 11 h of ambient solar radiation (from 0700 HR to 1800 HR) and 8 h of supplemental metal halide 7500 °K lamps (UltraSun 1000 W; Hawthorne Hydroponics LLC., Vancouver, WA, United States) that operated from 1700 HR to 0100 HR. Greenhouse environmental conditions were recorded every 15 min by a weather station data logger (WatchDog 2475; Spectrum Technologies, Inc., Aurora, IL, United States). Average air temperature, relative humidity, photosynthetic active radiation, and DLI was 24.0 ± 0.08 □, 60.1 ± 0.45 %, 261.6 ± 5.07 µmol·m^-2^·s^-1^, and 22.6 ± 0.44 mol·m^-2^·d^-1^, respectively.

After transplant in 1.1 L containers, all plants were cultivated in seven identical environmentally controlled rooms, as described previously. Average air temperature, relative humidity, and light intensity for the vegetative stage and flowering stage for each lighting treatment were also reported in Supplementary table 2.

#### Plant measurements and data collection

Plant height, recorded from the substrate surface to the tallest meristem, was measured at initiation of lighting treatments and at flowering. Days to flower and plant sex were recorded when plants started to flower, as defined previously. Flowering of monecious plants was defined when female or male flowering occurred as defined previously or by decimal code of 2301 and 2304, respectively (Mediavilla et al., 1998). Boolean evaluation of plant flowering status was conducted at the end of week 5 after the initiation of the lighting treatments. Experimental design and data analysis were conducted as described for Expt. 1.

### Expt.3: Expanded photoperiod trial for selected essential oil and fiber/grain cultivars

Based on results of Expt. 1 and 2., an expanded photoperiod trial was designed with select essential oil, fiber, and grain cultivars to better understand the effect of photoperiodism on a broader scale.

#### Seedling preparation and vegetative stage

Six fiber/grain hemp cultivars, ‘Carmagnola Selezionata’, ‘Helena’, ‘Eletta Campana’, ‘HAN-FN-H’, ‘PUMA-3’, and ‘PUMA-4’, were propagated as described in Expt.2 on May 24, 2020. Cuttings of 10 essential oil cultivars, ‘ACDC-AC’, ‘Super CBD-AC’, ‘Cherry-AC’, ‘Cherry Blossom-BC’, ‘Cherry Wine-BS’, ‘Cherry Blossom-BS’, ‘Cherry*T1-BS’, ‘JL Baux-CC’, ‘ACDC-CC’, and ‘Cherry-CC’, were propagated as described in Expt. 1 on June 18, 2020. Both cuttings and seeded trays were placed under a mist bench that misted 8 s every 20 min in a greenhouse and grown at 25 □ under natural daylight with supplemental metal halide lamps as described in Expt. 2 maintaining an 18 h photoperiod. Plants grew vegetatively under the mist bench in the greenhouse for 3-4 weeks before being transplanted into 1.1 L containers and assigned to lighting treatments.

Seedlings or clones of the 10 essential oil cultivars were thinned and transplanted as described in Expt. 2 with Pro-Mix HP soilless substrate on June 18, 2020, for fiber cultivars, and July 11, 2020, for essential oil cultivars. Slow-release fertilizer was applied as described in Expt. 1. All plants were cultivated for vegetative growth for 7 d and then randomly assigned to identical environmentally controlled rooms under different lighting treatments with 5 replicates per each cultivar.

#### Lighting treatments during flowering stage

Six lighting treatments were randomly assigned to each controlled room as proposed: 12 h 30 min (0600 HR to 1830 HR), 13 h (0600 HR to 1900 HR), 14 h 30 min (0600 HR to 2030 HR), 14 h 45 min (0600 HR to 2045 HR), 15 h (0600 HR to 2100 HR), and 15 h 30 min (0600 HR to 2130 HR). Different photoperiods were provided by LEDs as described previously. Photoperiod lighting treatments were maintained for five weeks before the termination of the experiment. Ten essential oil cultivars were selected based on the results from Expt. 1 and were evaluated from 14 h 30 min to 15 h 30 min.

A Boolean evaluation of flowering status was conducted as described previously. Environmental conditions of the greenhouse during the vegetative stage were as described in Expt. 2 and the environmental conditions of the controlled rooms during the flowering stage were as described in Expt. 1 and provided in Supplementary table 2. Experimental design and data analysis were conducted as described for Expt. 1.

### Expt.4: Flowering time trial under natural daylengths within a field-grown subtropical Central Florida environment

#### Seedling preparation and vegetative stage

Fourteen essential oil and 18 fiber/grain cultivars were evaluated for flowering response time under natural daylength, field-grown conditions following seedling establishment of fiber/grain cultivars and rooting of clonally propagated essential oil cultivars. Fiber/grain seeds were sown in 72-cell trays within Pro-Mix HP soilless substrate on April 30, 2020 and propagated as described in Expt. 2. Seedlings were watered daily by hand as needed. Fourteen essential oil cultivars were clonally propagated as described in Expt. 2 on May 1, 2020. Rooted plants were transplanted into field on June 3, 2020.

#### Field trial set up

The field trial was designed using plasticulture production techniques with Chapin Turbulent Flow-Deluxe drip tape (Catalog # 11714142N, Jain Irrigation USA, Watertown, NY, USA) placed below the plastic emitting 0.76 L h^-1^ per dripper at 68.9 kPa spaced 0.10 m between drippers. Plants were spaced 0.9 m apart within rows and rows were spaced 1.5 m apart between row centers. Total plot lengths were 3.7 m including walking allies. Total trial area was 0.9 ha. Trials received 2 hours of drip irrigation per day. A soluble fertilizer with micronutrients (Peter 20-20-20; ICL Specialty Fertilizers, Dorchester County, SC, U.S.), was applied every 14 d at a rate of 8.8 kg N ha^-1^ for an accumulated rate of 48 kg N ha^-1^ (6 applications total).

#### Experimental design and data collection

The experiment was conducted using a complete randomized block design comprised of an essential oil trial and a fiber/grain trial. Both trials contain three replicates of each cultivar. Each plot/replicate within the trial consisted of three plants. Flowering time was measured as defined previously. Light intensity during civil twilight period (sun 6 to 0 degrees below horizon) was recorded every two minutes manually in an open field with a quantum sensor (MQ-500; Apogee Instruments Inc., Logan, UT, United States) for three days. A restricted maximum likelihood mixed model analysis in JMP^®^ Pro 15 (SAS Institute, Inc., Cary, NC, United States) was performed to estimate genetic means of flowering time.

## RESULTS AND DISCUSSIONS

### Identifying Critical Photoperiod Thresholds

Critical photoperiod differed significantly among essential oil cultivars (Table 1). In addition, a significant effect was observed on flowering percentages of the essential oil cultivars. As expected, all essential oil cultivars flowered in response to 12 h photoperiod and no plants flowered in response to 18 h (Table 1). One cultivar, ‘Cherry Wine-CC’, was identified with a critical photoperiod below 14 h, with no flowers developed under 14 h. Four cultivars expressed 100% floral initiation at the longest photoperiod (14 h, excluding 18 h control) with an additional six cultivars that demonstrated a complete floral initiation (>50%) when cultivated under 14 h of light. For this reason, expanded photoperiod treatments of up to 15 h 30 min were evaluated for select essential oil cultivars (Expt. 3). Of the 10 essential oil cultivars evaluated within Expt. 3, five cultivars expressed a majority (>50%) of floral initiation between 15 h and 15 h 30min. Four of them, including ‘Cherry-AC’, ‘Cherry Blossom-BS’, ‘ACDC-CC’, and ‘Cherry-CC’, have been identified with a critical photoperiod within this range. In addition, ‘Cherry Wine-CC’ had the shortest, critical photoperiod identified between 13 h 45 min and 14 h. The critical photoperiod for ‘Super CBD-AC’ and ‘Cherry Blossom-BC’ occurred between 14 h 45 min and 15 h. For the rest of the cultivars, ‘ACDC-AC’ had a significant flowering reduction when the photoperiod was extended from 15 h to 15 h 30min. ‘Wife-AC’ flowered significantly less under 13 h 30 min compared to 13 h, but the critical photoperiod is likely greater than 14 h. Similarly, the percentage flowering of ‘Berry Blossom-BS’, ‘Cherry Blossom-Tuan-BS’, and ‘Wife-CC’ decreased when photoperiod was increased from 13 h 45 min to 14 h. Our results suggest that a photoperiod difference as little as 15 min could have a significant influence on floral initiation and development of some essential oil hemp cultivars. Moreover, floral initiation can occur at varying rates when the photoperiod is close to the critical threshold of some cultivars.

**Table 1.**
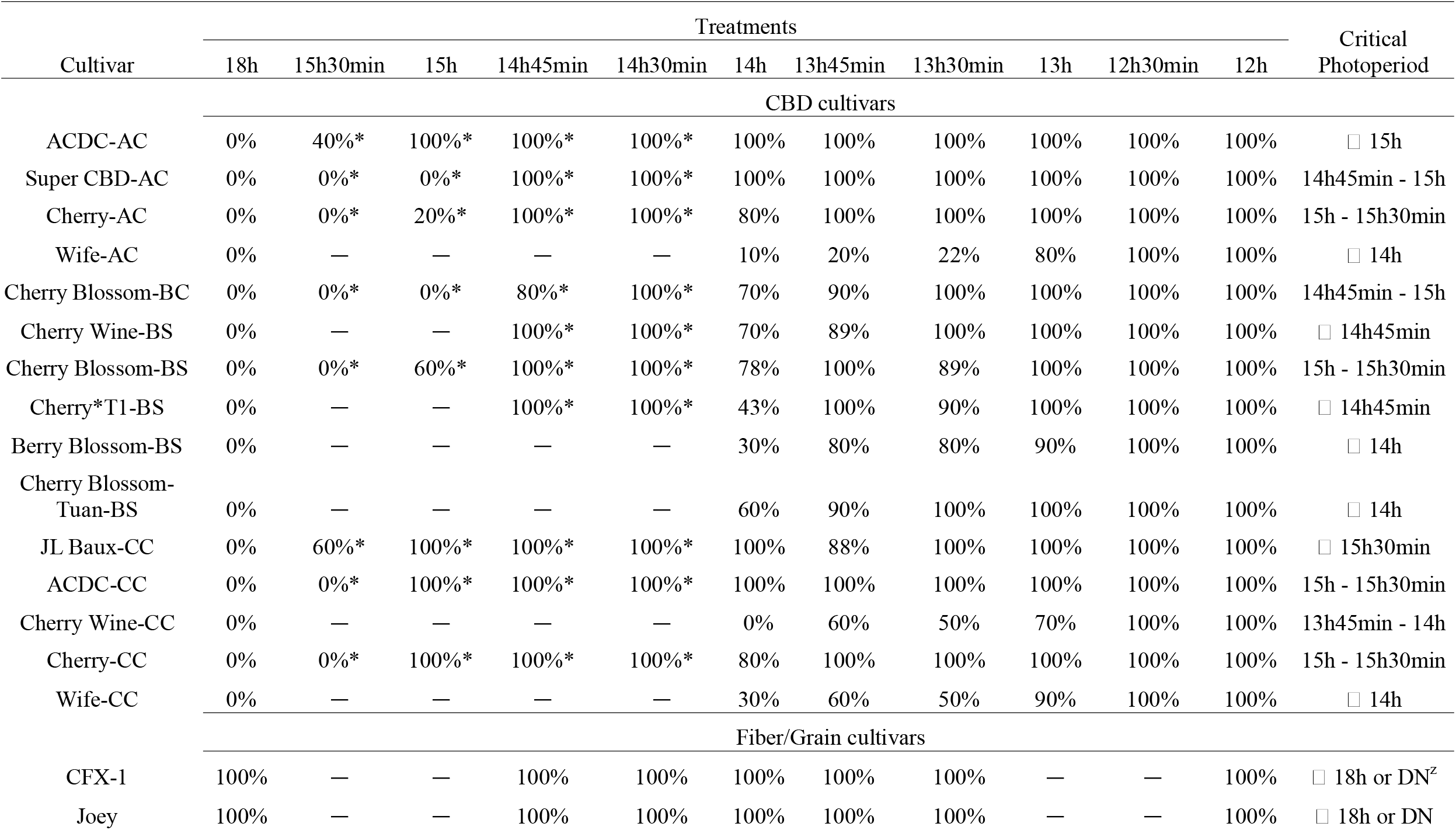
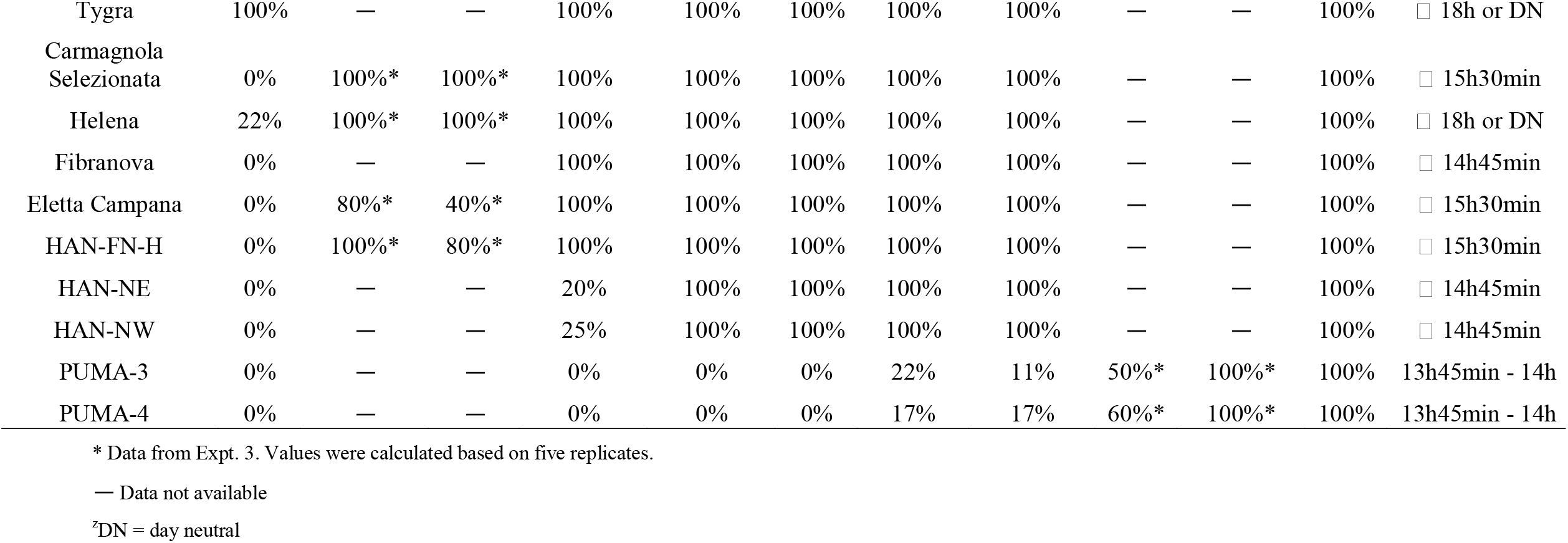
Flowering percentage of CBD and fiber hemp from Expt. 1, 2 and 3. The Boolean evaluation was conducted at week 5 of flowering after the treatment initiation excluding dead plants.

| Cultivar | Treatments |  |  |  |  |  |  |  |  |  |  | Critical<br>Photoperiod |
| --- | --- | --- | --- | --- | --- | --- | --- | --- | --- | --- | --- | --- |
|  | 18h | 15h30min | 15h | 14h45min | 14h30min | 14h | 13h45min | 13h30min | 13h | 12h30min | 12h |  |
|  | CBD cultivars |  |  |  |  |  |  |  |  |  |  |  |
| ACDC-AC | 0% | 40% * | 100% * | 100% * | 100% * | 100% | 100% | 100% | 100% | 100% | 100% | ☐ 15h |
| Super CBD-AC | 0% | 0% * | 0% * | 100% * | 100% * | 100% | 100% | 100% | 100% | 100% | 100% | 14h45min - 15h |
| Cherry-AC | 0% | 0% * | 20% * | 100% * | 100% * | 80% | 100% | 100% | 100% | 100% | 100% | 15h - 15h30min |
| Wife-AC | 0% | — | — | — | — | 10% | 20% | 22% | 80% | 100% | 100% | ☐ 14h |
| Cherry Blossom-BC | 0% | 0% * | 0% * | 80% * | 100% * | 70% | 90% | 100% | 100% | 100% | 100% | 14h45min - 15h |
| Cherry Wine-BS | 0% | — | — | 100% * | 100% * | 70% | 89% | 100% | 100% | 100% | 100% | ☐ 14h45min |
| Cherry Blossom-BS | 0% | 0% * | 60% * | 100% * | 100% * | 78% | 100% | 89% | 100% | 100% | 100% | 15h - 15h30min |
| Cherry*T1-BS | 0% | — | — | 100% * | 100% * | 43% | 100% | 90% | 100% | 100% | 100% | ☐ 14h45min |
| Berry Blossom-BS | 0% | — | — | — | — | 30% | 80% | 80% | 90% | 100% | 100% | ☐ 14h |
| Cherry Blossom-Tuan-BS | 0% | — | — | — | — | 60% | 90% | 100% | 100% | 100% | 100% | ☐ 14h |
| JL Baux-CC | 0% | 60% * | 100% * | 100% * | 100% * | 100% | 88% | 100% | 100% | 100% | 100% | ☐ 15h30min |
| ACDC-CC | 0% | 0% * | 100% * | 100% * | 100% * | 100% | 100% | 100% | 100% | 100% | 100% | 15h - 15h30min |
| Cherry Wine-CC | 0% | — | — | — | — | 0% | 60% | 50% | 70% | 100% | 100% | 13h45min - 14h |
| Cherry-CC | 0% | 0% * | 100% * | 100% * | 100% * | 80% | 100% | 100% | 100% | 100% | 100% | 15h - 15h30min |
| Wife-CC | 0% | — | — | — | — | 30% | 60% | 50% | 90% | 100% | 100% | ☐ 14h |
|  | Fiber/Grain cultivars |  |  |  |  |  |  |  |  |  |  |  |
| CFX-1 | 100% | — | — | 100% | 100% | 100% | 100% | 100% | — | — | 100% | ☐ 18h or DN<sup>z</sup> |
| Joey | 100% | — | — | 100% | 100% | 100% | 100% | 100% | — | — | 100% | ☐ 18h or DN |
| Tygra | 100% | — | — | 100% | 100% | 100% | 100% | 100% | — | — | 100% | □ 18h or DN |
| Carmagnola<br>Selezionata | 0% | 100%* | 100%* | 100% | 100% | 100% | 100% | 100% | — | — | 100% | □ 15h30min |
| Helena | 22% | 100%* | 100%* | 100% | 100% | 100% | 100% | 100% | — | — | 100% | □ 18h or DN |
| Fibranova | 0% | — | — | 100% | 100% | 100% | 100% | 100% | — | — | 100% | □ 14h45min |
| Eletta Campana | 0% | 80%* | 40%* | 100% | 100% | 100% | 100% | 100% | — | — | 100% | □ 15h30min |
| HAN-FN-H | 0% | 100%* | 80%* | 100% | 100% | 100% | 100% | 100% | — | — | 100% | □ 15h30min |
| HAN-NE | 0% | — | — | 20% | 100% | 100% | 100% | 100% | — | — | 100% | □ 14h45min |
| HAN-NW | 0% | — | — | 25% | 100% | 100% | 100% | 100% | — | — | 100% | □ 14h45min |
| PUMA-3 | 0% | — | — | 0% | 0% | 0% | 22% | 11% | 50%* | 100%* | 100% | 13h45min - 14h |
| PUMA-4 | 0% | — | — | 0% | 0% | 0% | 17% | 17% | 60%* | 100%* | 100% | 13h45min - 14h |

\* Data from Expt. 3. Values were calculated based on five replicates.
— Data not available
<sup>z</sup>DN = day neutral

Less variation in critical photoperiod thresholds were observed for fiber/grain hemp than essential oil cultivars. Similar to essential oil cultivars, all fiber/grain cultivars flowered in response to 12 h photoperiod (Table 1). For the majority of the fiber cultivars (8 of 12), plants did not flower under an 18 h photoperiod. ‘CFX-1’, ‘Joey’, ‘Tygra’, and ‘Helena’ flowered in response to a photoperiod of 18 h and did not remain vegetative like the majority of the other fiber hemp cultivars evaluated in this study, thus suggesting their critical photoperiod could be above 18 h. The critical photoperiod of PUMA 3 and 4 was identified between 13 h 45 min and 14 h, but the floral initiation was greatly reduced by more than 70% when day length exceeded 13 h. Similarly, the flowering of HAN-NE and HAN-NW was also greatly reduced when day length exceeded 14 h 30 min. To verify the critical photoperiod of ‘CFX-1’, ‘Joey’, ‘Tygra’, and ‘Helena’, seeds were germinated on February 18, 2020, and placed under 24 h photoperiod in a greenhouse. All four cultivars flowered on April 20, 2020 under a 24 h photoperiod. This observation was consistent with previous reports where 24 h photoperiod did not prevent the flowering of ‘Fedrina 74’ and ‘Kompolti Hybrid TC’ and that the critical photoperiod of a Portuguese fiber hemp variety from Coimbra is between 20 and 24 h (Heslop-Harrison and Heslop-Harrison, 1972; Van der Werf et al., 1994). In addition, ‘CFX-1’ and ‘Joey’ formed flower buds during the three-week propagation stage in the greenhouse in Expt. 2. Available literature supports that primordium formation in hemp varieties occurs in response to quantitative short days and the photoperiod inductive phase is jointly affected by photoperiod and temperature (Lisson et al., 2000; Hall et al., 2012). However, Spitzer-Rimon et al. (2019) argued that *Cannabis* can enter the reproductive phase under both long-day and short-day conditions because “solitary flowers”, which being developed in the axil of each stipulate leaf, are differentiated under such conditions, and flower induction of “solitary flowers” is likely age-dependent and is controlled not by photoperiod, but rather internal signals. Therefore, they reported that *Cannabis* can be considered a day-neutral plant where floral initiation is not dependent upon photoperiod requirements. These “solitary flowers”, which can be referred to as sex-indicating flowers or pre-flowers, are believed to be the start of calyx development in hemp and are not photoperiod dependent (Green 2017; Williams, 2020). In our study, long-day conditions did not prevent the floral initiation of ‘CFX-1’, ‘Joey’, ‘Tygra’, and ‘Helena’. Thus, they are likely day-neutral cultivars given floral initiation occurred in response to 24 h photoperiod.

Hall et al. (2012) suggested that the hemp juvenile phase was not affected by photoperiod and the length of the juvenile phase is either determined by development of reproductive organs or the apical meristem, which is independently timed to produce flowering signals. In our study, the length of juvenile phase was observed to be cultivar specific, with ‘CFX-1’ being the shortest and ‘Helena’ being the longest among the four, day-neutral cultivars (Fig 2). Traits, such as days to maturity and cannabinoid production, have been identified to be nearly entirely controlled by genetics. However, environment can play a significant role for other traits, such as yield and plant height, and thus the influence of environment and genetics are likely needed to be considered collectively (Campbell et al., 2019). Different hemp cultivars have been suggested to have different lengths of juvenile phase and photo-sensitive phase largely in association with geographic origin. Cultivars adapted to northern latitudes tend to have a short life cycle and grow and flower faster within their limited growing seasons, whereas cultivars adapted to southern latitudes and closer to the equator tend to flower later to ensure sufficient vegetative growth before short days occur (Amaducci et al., 2008; Small, 2015; Zhang et al., 2018). This theory is supported by our study results where cultivars of northern origin (‘CFX’, ‘Joey’, ‘Tygra’, etc.) responded to a longer photoperiod and flowered faster than southern cultivars (‘PUMA-3’, ‘PUMA-4’, ‘HAN-NW’, etc.) having a shorter critical day length threshold (Fig 2). Thus, understanding the juvenile phase and photosensitivity is essential for selecting the right hemp cultivar for a target region.

**Figure 1.**
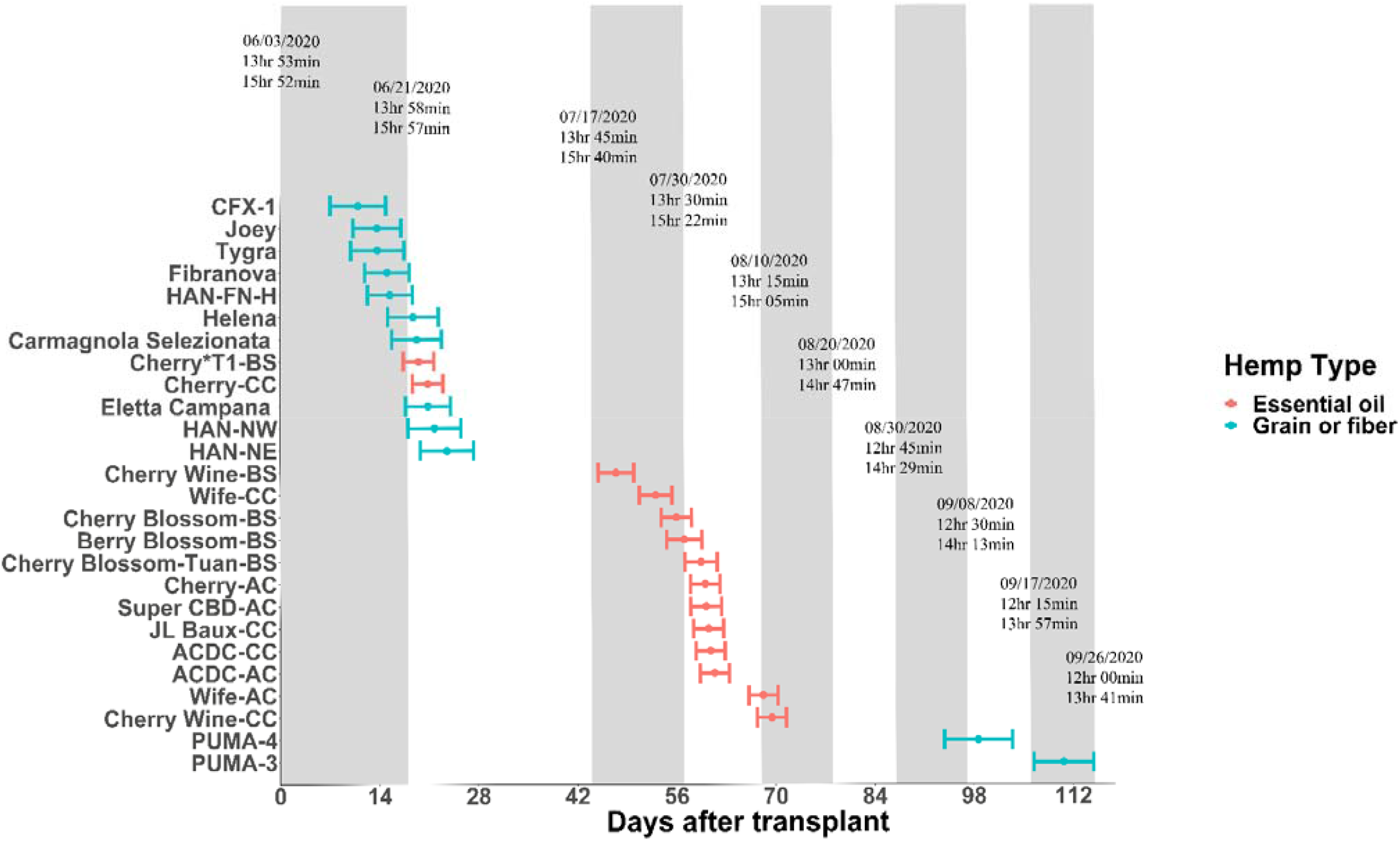
Flowering date of essential oil, grain, and fiber cultivars after transplanting from greenhouse to field conditions on June 3^rd^, 2020. Points depict genetic means and error bars represent 95% confidence intervals. Floating text depicts date of 15 min daylength intervals (top), sunrise to sunset daylengths (middle), and civil twilight lengths (bottom).

**Figure 2.**
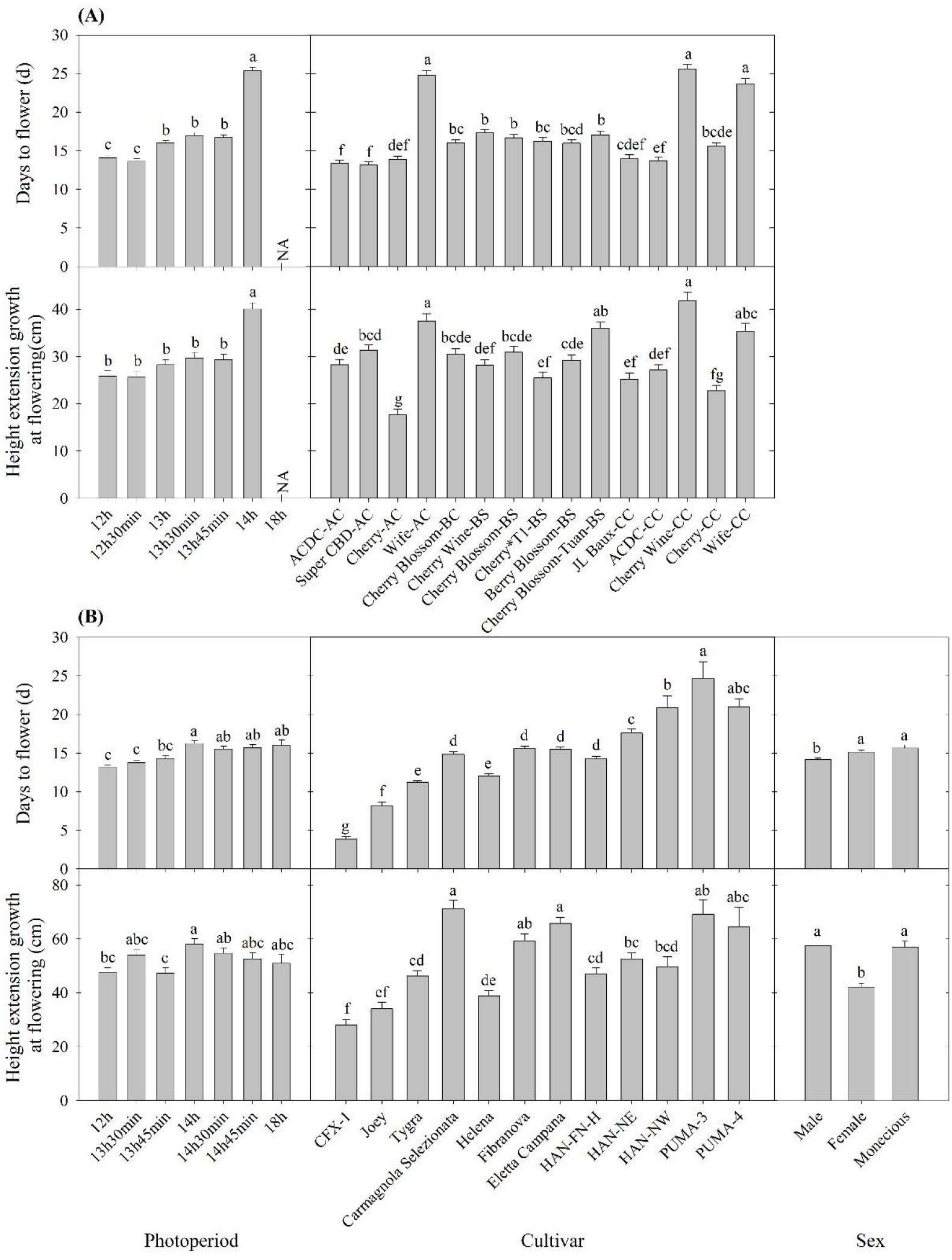
Days to flower after lighting treatment initiated and height extension growth at flowering of (A) essential oil and (B) fiber/grain cultivars under different photoperiods, cultivars, and/or sex from Expt. 1 and 2. Means sharing a letter are not statistically different by Tukey’s HSD test at P ≤ 0.05. Error bars indicate standard error.

Some plant species can respond to light even at a very low intensity and are thus considered highly photosensitive. For these species of plants, civil twilight, or the period of time that occurs shortly before sunrise and after sunset when the sun is between 0 and 6 degrees below the horizon, may still be biologically effective to the plant’s photoperiodism response (Kishida, 1989; Salisbury, 1981). For example, rice (Oryza sativa) is light-insensitive to twilight both at dusk and dawn; perilla (Perilla frutescens) and Biloxi soybean (Glycine max) are light-insensitive at dusk but more light-sensitive at dawn; and cocklebur (Xanthium saccharatum) is both light-sensitive at dusk and dawn (Takimoto and Ikeda, 1961). For hemp, Borthwick and Scully (1954) suggested that 0.12 foot-candle or more would sufficiently prevent hemp from flowering, suggesting hemp is rather sensitive to light. In our experience, light intensity as little as 2 µmol·m^-2^·s^-1^ can cause light pollution and disrupt vegetative growth and transition hemp into flowering for ‘Cherry Blossom-BS’ in the greenhouse. To evaluate the effect of twilight on hemp, we conducted field trials (Expt. 4) to investigate the performance of essential oil, fiber, and grain hemp cultivars under natural day light conditions. The average light intensity during civil twilight period in our study was 2.4 ± 0.54 µmol·m^-2^·s^-1^, which is within the light range reported by Kishida (1989). By comparing plant response and photoperiod to sunrise to sunset daylengths and civil twilight lengths, we concluded that flowering performance of hemp is affected by civil twilight (Fig. 1). Most flowering data from the field trial was in alignment with our trials from the controlled rooms (Expt. 1 to 3) and plants flowered within the critical photoperiod we tested, with only a few exceptions. ‘JL Baux-CC’ and ‘ACDC-AC’ flowered later and slower in the field compared to the controlled rooms (Fig. 1). It is possible that the day length changes under the natural conditions are slower to occur and not as drastic compared to conditions imposed in the controlled rooms and therefore plants would respond to day length changes slower under natural conditions. On the contrary, ‘Super CBD-AC’ and ‘Cherry-CC’ flowered earlier, suggesting that they might be more sensitive to the dark period. In addition, differences in individuals perception of flowering initiation may have led to the reduced correlation between field and growth chamber floral initiation dates collected for these cultivars, additional years of field trials will aid in the importance of civil twilight’s effect on *Cannabis* flowering. Collectively, we believe that civil twilight length and the slow progression of day length changes under natural conditions should be taken into consideration for the biologically effective photoperiod for hemp flowering.

### Days to Flower

Flowering response was delayed as flowering photoperiod increased. In both essential oil and fiber/grain cultivars, plants subjected to 12 h photoperiod had an average flowering time of 13-14 d (Fig. 2). This is supported by Borthwick and Scully (1954) where 10-14 d of short-day photoperiod was sufficient for flower induction in at least some of the Chilean and Kentucky varieties.

Among essential oil cultivars, flowering was generally delayed by 1 to 2 d when photoperiod exceeded 13 h compared to 12 h, and flowering was significantly delayed by 7 to 8 d when photoperiod exceed 14 h (Fig. 2). Across cultivars and regardless of sources, ‘ACDC’ and ‘Super CBD’ flowered the fastest, with an average flowering time of 13 d after initiation of the critical photoperiod. ‘Wife-AC’, ‘Wife-CC’, and ‘Cherry Wine’ had an average flowering time of 21 d, suggesting these cultivars took longer to either perceive the photoperiod or to complete flower formation and initiation.

Essential oil hemp cultivars demonstrated delayed floral initiation at longer photoperiods and significant genetic variation in floral initiation across photoperiod treatments. Variance in observed photosensitivity is likely a result of genetic variation that influenced floral initiation response to light cues. Flowering for most essential oil cultivars was delayed by 5 to 13 d under 14 h photoperiod compared to 13 h 45 min (Fig. 3 and Supplementary fig. 1). Flowering of ‘ACDC-AC’ and ‘Cherry*T1-BS’ was significantly delayed by 4 and 6 d, respectively, under 13 h 45 min compared to 12 h. Moreover, the delayed flowering of ‘Cherry-CC’ started at 13 h 30 min and in ‘Wife-CC’, 13 h. This suggests that floral initiation of these cultivars was more sensitive to photoperiod than others. In contrast, no significant differences were observed in days to flower among different treatments of ‘Wife-AC’, ‘Berry Blossom-BS’, and ‘Cherry Blossom-Tuan-BS’, suggesting the flowering formation and initiation were rather similar under different day lengths, as long as they were below the critical photoperiod.

**Figure 3.**
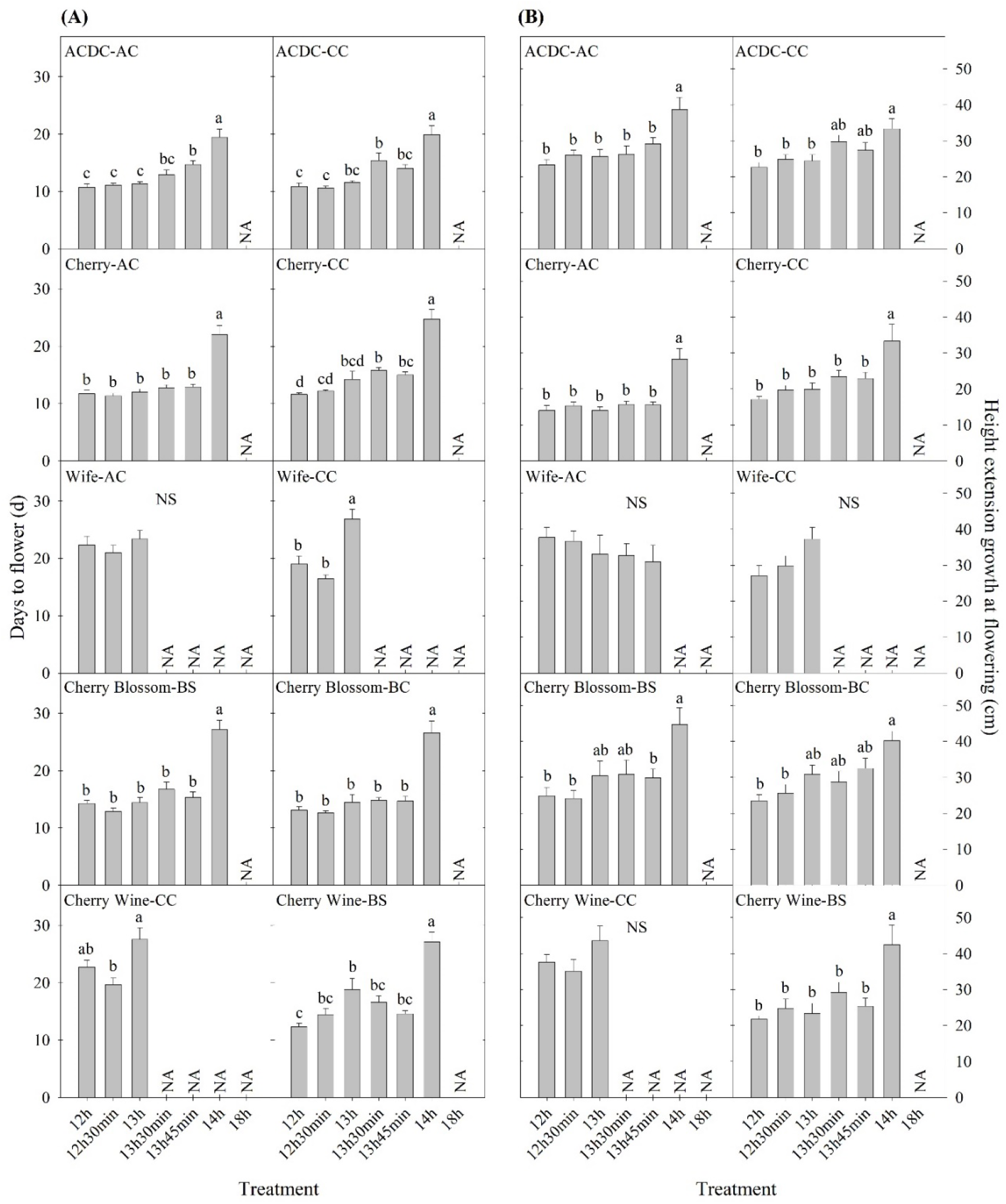
Comparison of (A) days to flower and (B) height extension growth at flowering of ten essential oil cultivars with same name but different sources in Expt. 1. All data were pooled from ten replications except ‘ACDC-AC’ (n=9). NS indicates insignificant treatment effects. NA indicates the majority of the 10 replicates (i.e. 6/10) were reported as not flowering. Means sharing a letter are not statistically different by Tukey’s HSD test at P ≤ 0.05. Error bars indicate standard error.

Cultivars represented by the same name acquired from different sources performed differently in days to flower. A photoperiod of 13 h 30 min significantly delayed the flowering of ‘ACDC-CC’ but not ‘ACDC-AC’, whereas 13 h 45 min delayed the flowering of ‘ACDC-AC’ but not ‘ACDC-CC’ (Fig. 3). Delay of flowering in ‘Cherry-CC’ started under a photoperiod of 13 h 30 min, while flowering occurred 30 min later at 14 h for ‘Cherry-AC’. A photoperiod of 13 h delayed flowering of ‘Wife-CC’ by 8 d compared to 12 h, but not in ‘Wife-AC’. Similarly, flowering of ‘Cherry Wine-BS’ was significantly delayed by 7 d under 13 h photoperiod compared to 12 h photoperiod, but no differences in flowering were observed in ‘Cherry Wine-CC’. These results, in conjunction with Sawler et al. (2015), indicated that plants with the same cultivar names from different sources could have varying genetics and subsequently performed differently.

Most fiber/grain cultivars tested did not flower under 18 h photoperiod (Table 1). Flowering was delayed by 1 to 3 d if the photoperiod exceeded 14 h and no differences were observed among treatments beyond 14 h (Fig. 2). This is consistent with the theory that hemp is a quantitative SDP and has a photoperiod of roughly 14 h where flowering would occur promptly below 14 h and flowering would be delayed under a longer photoperiod (Amaducci et al., 2008; Borthwick and Scully, 1954; Hall et al., 2014; Heslop-Harrison and Heslop-Harrison, 1969). When subjected to the critical photoperiod, average days to flower was shortest among cultivars from northern latitudes and longest among those from southern latitudes with a gradient response correlated to the cultivar’s genetic origin. More specifically, Canadian/Northern European cultivars flowered 4 to 11 d after lighting transition. Cultivars from comparatively lower latitudinal regions (Northern Chinese/Southern European) flowered from 12 to 16 d whereas Southern Chinese cultivars flowered 21 to 25 d (Fig. 2 and Supplementary fig. 2). Moreover, different photoperiod treatments did not affect the flowering of ‘CFX-1’, ‘Joey’, ‘Tygra’, ‘Carmagnola’, and ‘Helena’ (Supplementary fig. 2). Collectively, considering that 24 h day length did not prevent ‘CFX-1’, ‘Joey’, ‘Tygra’, and ‘Helena’ from flowering and their flowering process was not influenced by imposed photoperiods (Table 1 and Supplementary fig. 2), these four fiber cultivars are likely photoperiod insensitive or day-neutral.

Temperature differences and other stresses such as nutrient deficiencies can result in differences in flowering time (Amaducci et al., 2008; Hall et al., 2012). Amaducci et al. (2008) indicated that high temperature will accelerate flowering by decreasing the duration between the formation of flower primordia and full flowering, and thus modeling had been used to predict flowering time based on day length and temperature. Hall et al. (2012) also indicated that the photoperiod inductive phase of hemp is jointly influenced by both air temperature and photoperiod. In our study, growing conditions including air temperature and nutrient fertility are nearly identical in each environmentally controlled room and thus did not contribute to differences in plant flowering performances. Flowering response, as recorded in this study, was therefore free from the confounding influence of temperature and nutrient deficiency (excluding Expt. 4) and thus provides an enhanced foundational understanding of relationships between photoperiod and flowering response in hemp.

Collectively, based upon study findings and available literature, we believe the hemp juvenile phase to be controlled by genetics rather than photoperiod or temperature. The pre-flowering of the single sex-indicating flower at the axillary is photo insensitive. The response to photoperiod from pre-flowering to flowering at the apical meristem is affected by both photoperiod and temperature and can be either quantitative (most cultivars) or day-neutral (such as ‘CFX-1’, ‘Joey’, ‘Tygra’, and ‘Helena’), dependent upon cultivar.

### Extension Growth

Generally, plant height increased as day length increased as would be expected from increased photosynthesis. Plant height extension growth was 47 to 102% greater under 14 h day length compared to the 12 h, control group in nine essential oil cultivars, while the imposed photoperiods did not affect the remaining six cultivars (Fig. 2). The longer stem could have resulted from a longer vegetative stage caused by the delay in floral initiation, which has been reported on a variety of crops (Craig and Runkle, 2013; Zhang and Runkle, 2019). Under certain photoperiod treatments, flowering was delayed but the height extension growth was not affected; this included ‘Cherry Wine-BS’, ‘Cherry Wine-CC’, and ‘Wife-CC’ under 13 h, ‘ACDC-CC’ under 13 h 30 min, ‘ACDC-AC’ and ‘Cherry*T1-BS’ under 13 h 45 min, and ‘Cherry-CC’ under both 13 h 30 min and 13 h 45 min (Fig. 3 and Supplementary fig. 1). Results indicated flowering initiation of essential oil hemp was more sensitive than extension growth in response to photoperiods. In addition, unlike flowering, extension growth of essential oil cultivars with the same name but from different sources generally responded similarly, except “Cherry Wine”. Campbell et al. (2019) indicated that plant height was collectively influenced by both environment (e.g. irrigation) and genetics, accounting for 38% and 36% of variance, respectively. We concluded that similar height extension growth among essential oil cultivars under different treatments was due to similar irrigation applications.

In contrast to essential oil cultivars, extension growth of fiber/grain cultivars was not affected by photoperiod, except ‘CFX-1’ and ‘HAN-NE’ (Supplementary Fig. 3). Height extension of ‘HAN-NE’ was 58 to 64% greater when day length exceeds 14 h due to the later flower initiation development. Interestingly, stem extension of ‘CFX-1’ was shorter under 18 h photoperiod compared to 13 h 30 min. We believe ‘CFX-1’ is photo insensitive and this difference was caused by individual variances.

### Sex

Plant sex was recorded and calculated across the lighting treatments for fiber/grain hemp cultivars (Table 2). Among all the fiber/grain hemp cultivars tested, most cultivars had a relatively equal proportion of male and female plants in general with a small occurrence of monecious plant development, which is consistent with Hall et al. (2012). ‘Tygra’ had the highest proportion of monecious plants (55.7%) with a small proportion of female plant development (4.3%). ‘Helena’ was one-third monecious. Most of ‘CFX-1’ were female plants with no monecious plants that developed. The sex of more than 25% of ‘HAN-NE’, ‘HAN-NW’, ‘PUMA-3’, and ‘PUMA-4’ could not be determined due to a lack of flowering response throughout the duration of our experiment.

**Table 2.** Origin and sex status of fiber/grain hemp cultivars in Expt. 2.

| Cultivar | Origins | Male | Female | Monecious | Unknown |
| --- | --- | --- | --- | --- | --- |
| CFX-1 | Canadian | 17.1% | 82.9% | 0.0% | 0.0% |
| Joey | Canadian | 45.7% | 34.8% | 19.5% | 0.0% |
| Tygra | Poland | 40.0% | 4.3% | 55.7% | 0.0% |
| Carmagnola Selezionata | Southern European | 48.1% | 33.3% | 3.7% | 14.9% |
| Helena | Southern European | 50.7% | 10.2% | 33.3% | 5.8% |
| Fibranova | Southern European | 32.7% | 46.9% | 6.1% | 14.3% |
| Eletta Campana | Southern European | 35.7% | 47.1% | 2.9% | 14.3% |
| HAN-FN-H | Northern Chinese | 32.8% | 48.6% | 4.3% | 14.3% |
| HAN-NE | Central Chinese | 37.1% | 37.1% | 0.0% | 25.8% |
| HAN-NW | Central Chinese | 25.0% | 50.0% | 0.0% | 25.0% |
| PUMA-3 | Southern Chinese | 14.3% | 7.9% | 1.6% | 76.2% |
| PUMA-4 | Southern Chinese | 7.3% | 12.2% | 0.0% | 80.5% |

Overall, flowering of female and monecious plants was delayed by 1-2 d compared to male plants (Fig. 3). Our observations were supported by Borthwick and Scully’s (1954) findings where greater flowering delay occurred in male plants when compared to female plants under long photoperiods. Hall et al. (2012) and van der Werf et al. (1994) suggested that extending day length would alter the sex proportion of flowering hemp plants and that male and monoecious plants would fail to flower when photoperiod exceeded the optimal day length. Additionally, Werf et al. (1994) suggested that unlike male hemp plants, female flowering would be less influenced by photoperiod. We did not observe such trends in fiber hemp cultivars and no flowering pattern or changes in flowering percentage were identified when photoperiod exceeded the optimal 14 h (Supplementary Table). Female hemp also has a significantly shorter extension growth than male and monecious plants at flowering (Fig. 3).

## CONCLUSION

This research reported flowering and growth of 27 hemp cultivars in response to different photoperiods under both indoor controlled and outdoor natural environments. Pre-flowering of hemp is photo insensitive, but the response to photoperiod from pre-flowering to flowering can be either quantitative or day neutral. Depending on photosensitivity, a photoperiod difference as little as 15 min significantly influenced floral initiation of some essential oil cultivars. Northern fiber/grain hemp cultivars had a shorter juvenile phase and faster flowering than cultivars from southern latitudes. Cultivar name may not be enough to finely estimate photoperiod response for essential oil cultivars. Flowering performance of hemp appears to be influenced by civil twilight and thus this should be a consideration when attempting to time cultivation to maximize vegetative and flowering response. Male plants flower faster than female and monecious plants. Plant height generally increased as the day length increased in essential oil cultivars but not in fiber/grain cultivars.

## SUPPORTING INFORMATION

**S1 Table**. Germplasm source, types, and referred name for fifteen essential oil hemp cultivars.

**S2 Table**. Means (± SE) of controlled rooms air temperature, relative humidity, and photosynthetic photon flux density (PPFD) of photoperiod treatments for essential oil, fiber, and grain cultivars in Expt. 1, 2, 3, and 4 as measured by thermocouples and quantum sensors. Light intensity was measure at ten representative positions at plant canopy level at the onset of each growing stage while air temperature and relative humidity were recorded by a wireless data logger throughout the growing stages.

**S1 Figure**. Days to flower (A) and height extension growth at flowering (B) of five essential oil cultivars in Expt. 1. All data were pooled from ten replications except ‘JL Baux-CC’ (eight replications). NS indicates insignificant treatment effects. NA indicates the majority of the 10 replicates (i.e. 6/10) were reported as not flowering. Means sharing a letter are not statistically different by Tukey’s HSD test at P ≤ 0.05. Error bars indicate standard error.

**S2 Figure**. Days to flower of twelve fiber/grain hemp cultivars in Expt. 3. All data were pooled from multiple replications as described in the Methods. NS indicates insignificant treatment effects. NA indicates the less than three plants were reported as not flowering. Means sharing a letter are not statistically different by Tukey’s HSD test at P ≤ 0.05. Error bars indicate standard error.

**S3 Figure**. Height extension growth at flowering of twelve fiber/grain hemp cultivars in Expt. 3. All data were pooled from multiple replications as described in the Methods. NS indicates insignificant treatment effects. NA indicates the less than three plants were reported as not flowering. Means sharing a letter are not statistically different by Tukey’s HSD test at P ≤ 0.05. Error bars indicate standard error.

## ACKNOWLEDGEMENT

We would like to acknowledge Brandon White and Chris Halliday for their technical support; James Johnston and Dillan Raab for their hard work and effort maintaining experimental plants and collecting phenotypic data; Jerry Fankhauser and Sandra Alomar for administrative assistance; Green Point Research, ANO CBD, and Green Roads for donating the cultivars used in this research; and all members of the University of Florida IFAS Industrial Hemp Pilot Project for their collaboration.

## AUTHOR CONTRIBUTIONS

MZ: conceptualization, data curation, formal analysis, investigation, methodology, original draft, review and editing (lead), supervision, validation, visualization.

SA: conceptualization, data curation, formal analysis, investigation, methodology, supervision, validation, visualization, review and editing (supporting).

ZB: funding acquisition, project administration, supervision, resources, review and editing (supporting).

BP: conceptualization, methodology, project administration, supervision, resources, review and editing (supporting).

